# Maternal cannabis use alters excitatory inputs to corticostriatal efferent neurons in rat offspring

**DOI:** 10.1101/2024.03.03.583210

**Authors:** Darren E. Ginder, Halle V. Weimar, Jonathan E.M. Lindberg, Zachary D.G. Fisher, Miranda M. Lim, James H. Peters, Ryan J. McLaughlin

## Abstract

With the recent surge in cannabis legalization across North America, there is legitimate concern that rates of cannabis use during pregnancy will dramatically increase in the coming years. However, the long-term impacts of prenatal cannabis exposure (PCE) on the brain and behavior remain poorly understood. Using a model of passive cannabis vapor exposure, we have previously shown that PCE impairs behavioral flexibility in an attentional set-shifting task in adult offspring, which is orchestrated in part by excitatory inputs from the medial prefrontal cortex (mPFC) to the nucleus accumbens (NAc). Given the fundamental role of these corticostriatal inputs in coordinating flexible reward-seeking strategies, we used a combination of retrograde tracing and *ex vivo* electrophysiology to test the hypothesis that maternal cannabis use alters the synaptic and intrinsic membrane properties of corticostriatal efferent neurons in exposed male and female rat offspring. Specifically, pregnant rat dams were trained to self-administer vaporized cannabis (69.7% THC; 150 mg/ml) twice daily throughout mating and gestation and offspring were subsequently injected with fluorescent retrobeads into the NAc core prior to conducting whole-cell *ex* vivo recordings of spontaneous excitatory and inhibitory post-synaptic currents (EPSC and IPSC, respectively) in retrolabeled mPFC neurons in adulthood. Our results indicate that PCE increases the frequency of spontaneous glutamatergic events (EPSCs) in NAc-projecting mPFC neurons in a sex-specific manner, which drives changes in excitatory to inhibitory (EPSC/IPSC) ratio, particularly in females. Furthermore, the amplitude of phasic glutamatergic events was reduced in cannabis-exposed offspring of both sexes, suggesting changes in postsynaptic receptor function. Altogether, these data demonstrate that PCE shifts the balance of excitatory/inhibitory inputs onto NAc-projecting mPFC neurons with limited effects on membrane conductance in females, resulting in reduced sex differences following maternal cannabis self-administration. These results provide putative neurophysiological mechanisms mediating previously observed behavioral changes, and future studies will need to test if these cannabis-induced changes are causal to long-term deficits in behavioral flexibility that have been previously documented in exposed offspring.

## 1. INTRODUCTION

The use of cannabis during pregnancy is a growing public health concern. Pregnant people are increasing their cannabis use following medical and recreational legalization (Young-Wolff et al., 2017) and these trends are expected to remain strong as the social stigma and perceived harms of cannabis use continue to decrease (Wen et al., 2018). A growing number of pregnant people believe that the use of cannabis during pregnancy is safer than the use of prescription medications (Foti et al., 2023), while during the COVID-19 pandemic, cannabis use among pregnant people significantly increased (Young-Wolff et al., 2021), predominantly to cope with mental health and stress-related issues (Young-Wolff et al., 2023). Furthermore, this is not solely a concern for the United States. Similar trends of increased cannabis use among pregnant people have been observed in Canada even prior to federal legalization, with an estimated 61% increase in the prevalence of maternal cannabis use from 2012 to 2017 (Corsi et al., 2019). Despite these concerning trends, the long-term effects of prenatal cannabis exposure (PCE) on brain development and function remain poorly understood. Thus, there is an urgent need for research that will increase our understanding of the long-lasting impact of PCE, particularly within brain circuits that are important for neurotypical cognitive and emotional functioning.

The primary psychoactive component of cannabis, Δ^9^-tetrahydrocannabinol (THC), is one of nearly 100 phytocannabinoids that have been identified in the cannabis plant (see Ashton, 2001 for review). THC is primarily responsible for most of the typical cannabimimetic effects of cannabis intoxication, which it accomplishes by binding the cannabinoid type-1 receptor (CB1R), a G-protein coupled receptor that is widely expressed on both excitatory and inhibitory presynaptic terminals (Maejima et al., 2001) in both humans (Glass & Faull, 1997) and rodents (Herkenham et al., 1990). CB1Rs are positively coupled to inwardly rectifying potassium channels (Pertwee, 1997) and maintain the resting membrane potential of the neuron by controlling potassium efflux (Howlett, 1995). Additionally, voltage gated Ca^2+^ channels are negatively coupled to CB1Rs (Mackie et al., 1995; Bayewitch et al., 1996), and their activation via either exogenous or endogenous cannabinoids ultimately decreases the probability of presynaptic neurotransmitter release. Thus, CB1R activity tightly constrains both excitatory and inhibitory synaptic neurotransmission. As such, perturbation within this system could have long-lasting consequences for the balance of excitation and inhibition within a given neural circuit.

A large body of literature has revealed a fundamental role for the endocannabinoid (eCB) system in regulating normal fetal neurodevelopment (see Pinky et al., 2019 for review). Autoradiography studies show that CB1Rs are highly expressed in human fetuses at an approximate gestational age of 20 weeks (Wang et al., 2003). Cell migration is under control of CB1R activity (see Harkany et al., 2008 for review) and exposure to cannabis *in utero* has been associated with increased cortical thickness (El Marroun et al., 2016). CB1R activity also controls axonal guidance and bundling (Berghuis et al., 2007; Fride et al., 2009) and synaptogenesis (Berghuis et al., 2007), all of which can be skewed following exposure to exogenous cannabinoids. THC readily crosses the placental barrier (Hutchings et al., 1989; Baglot et al., 2022), thereby directly impacting the developing fetus. Given that THC interacts primarily with the CB1R, the use of cannabis during pregnancy is likely to interfere with the ability of the eCB system to regulate the organization and function of synapses with high CB1R expression, thereby resulting in long-term alterations in cognition and behavior.

Accordingly, studies employing animal models of prenatal cannabinoid exposure have revealed long-term sex-dependent alterations in the function of mPFC neurons in exposed offspring. Prenatal exposure to THC or WIN 55,212-2 (WIN), a potent and selective CB1R agonist, alters long-term depression (LTD) and long-term potentiation (LTP) in layer V mPFC pyramidal neurons (Scheyer et al., 2020a; Scheyer et al., 2020b). Additionally, prenatal THC delays the switch of GABA channel polarity during early development from excitatory to inhibitory in the mPFC (Scheyer et al., 2020c). Moreover, prenatal WIN and prenatal THC exposure reduce LTD at excitatory synapses in the mPFC of male rats, but not females (Bara et al., 2018). Together, these studies indicate significant alterations to synaptic plasticity that skew the balance of excitation/inhibition in the mPFC to favor heightened excitability, particularly in male offspring exposed to cannabinoids *in utero*.

In line with the functional changes in mPFC neurons following prenatal cannabinoid exposure, human and rodent studies have revealed impairments in mPFC-dependent behaviors. Specifically, children exposed to cannabis *in utero* showed deficits in working memory (Fried et al., 1992; Leech et al., 1999) and difficulty with abstract thought, impulse control, and attention (Fried et al., 1998; Leech et al., 1999; Fried & Watkinson, 2001; El Marroun et al., 2011). Findings from our laboratory using a rat model of PCE corroborate these findings, showing a dysfunction of mPFC-dependent behavioral flexibility in offspring that lasts into adulthood (Weimar et al., 2020). Specifically, offspring non-contingently exposed to THC-dominant cannabis vapor showed deficits in their ability to shift strategies in an operant attentional set-shifting task when tested in adulthood (Weimar et al., 2020). Notably, this deficit has also been observed following selective inactivation of the mPFC-to-NAc pathway using an asymmetrical pharmacological disconnection approach (Block et al., 2007). Pharmacological inactivation of the mPFC on one side of the brain and inactivation of the NAc on the other side using bupivacaine resulted in similar deficits when rats were tested in a T-maze version of the same task, with rats requiring more trials to criterion to shift strategies (Block et al., 2007). Therefore, we reasoned that behavioral flexibility deficits previously reported by our lab (Weimar et al., 2020) may result from a cannabis-induced change in excitatory (or inhibitory) inputs to mPFC neurons that project to the NAc.

In the current study, we used a model of maternal cannabis use involving response-contingent delivery of vaporized cannabis extracts in pregnant rat dams to test whether PCE significantly alters the frequency and amplitude of spontaneous excitatory and inhibitory post-synaptic currents (EPSC and IPSC, respectively) in NAc-projecting mPFC neurons in adulthood. Furthermore, we examined differences between males and females on the above parameters. Based on the literature described above showing mPFC alterations exclusively in male offspring, we predicted that PCE in male offspring would increase the frequency of EPSCs in NAc-projecting mPFC neurons, while the frequency and amplitude of IPSCs would be reduced, resulting in a skewed EPSC/IPSC ratio that favors enhanced excitability of this pathway. On the other hand, due to the lack of sex differences seen in previous literature (Bara et al., 2018), we hypothesized that female cannabis-exposed offspring would experience no changes in any of the parameters tested.

## 2. METHODS

### 2.1 Animals

Rats were housed in a humidity-controlled animal suite on a 12 h on: 12 h off reverse light cycle with food and water available *ad libitum*. Nulliparous female Sprague Dawley rats (50 days old, Envigo, Indianapolis, IN) were pair-housed and randomly assigned to either the cannabis or vehicle (VEH) treatment group. Rats were weighed and handled daily. After one week of mating, males were removed and females were single housed until the day of birth, which was designated as postnatal day 0 (P0). Pups were cross fostered between P0 and P2 depending on the availability of litters. Before cross-fostering, each litter was weighed, and the number and sex of pups was determined. To distinguish prenatal treatment condition, a small ink dot was tattooed on the right or left hindpaw with Super Black^TM^ india ink (Speedball, Statesville, NC, USA). Litters were culled and subsequently cross fostered so that each dam raised approximately 10-12 pups. Ideally, each cross-fostered litter contained pups from both prenatal treatments; however, in some cases only one condition was raised by each dam due to differences in the timing of births between litters. All procedures were performed in accordance with the guidelines in the National Institute of Health Guide for the Care and Use of Laboratory Animals and were approved by the Washington State University Institutional Animal Care and Use Committee.

### 2.2 Drugs

Whole-plant cannabis extract was provided by the National Institute on Drug Abuse (NIDA) Drug Supply Program. According to the certificate of analyses provided upon shipment, the cannabis extract contained 69.81% THC, 0.89% tetrahydrocannabivarin (THCV), 0.83% cannabichromene (CBC), 2.69% cannabigerol (CBG), 1.51% cannabinol (CBN), and 0.73% Δ⁸ tetrahydrocannabinol (Δ⁸THC); cannabidiol (CBD) concentration was below the threshold of detection. The total terpene concentration was 1.35% (information regarding specific terpenes was not available). To prepare the final whole cannabis plant extract concentration, the raw cannabis plant extract was heated to 60℃ under constant stirring. The raw extract was suspended in 80% propylene glycol/20% vegetable glycerol (PG/VG) at a concentration of 150 mg/ml (mg extract/ml vehicle). This final concentration was selected based on previous studies from our lab comparing relative responding for cannabis vapor at different extract concentrations (Glodosky et al., 2020).

### 2.3 Maternal Cannabis Vapor Self-Administration

Maternal cannabis vapor self-administration followed previously described protocols (Weimar et al., 2023). In brief, vapor self-administration was conducted with a vapor delivery system running MED-Associates IV (Fairfax, VT) or La Jolla Alcohol Research Inc. software (LJARI; La Jolla, CA). Each vapor chamber (14.5” L x 10.5” W x 9.5” H) contains one active and one inactive nose-poke operandum (see **Figure 1A** for a schematic of the vapor chamber apparatus). Following a beam break in the active nose-poke, a signal was sent to the computer to release a 3 s puff of vapor and illumination of a cue light, followed by a 1 min timeout period. Nosepokes on the inactive operandum resulted in no vapor release or cue light illumination. Vapor was generated by a commercial e-cigarette SMOK Baby Beast TFV8 tank with a 0.20 M2 atomizer, 40-60 W range (SMOKtech, Shenzhen, China) containing the assigned VEH or cannabis solution. The system was under continuous vacuum, pulling vapor into the chamber and out through an exhaust port at an exchange rate of 1.0 L air/min.

**FIGURE 1:**
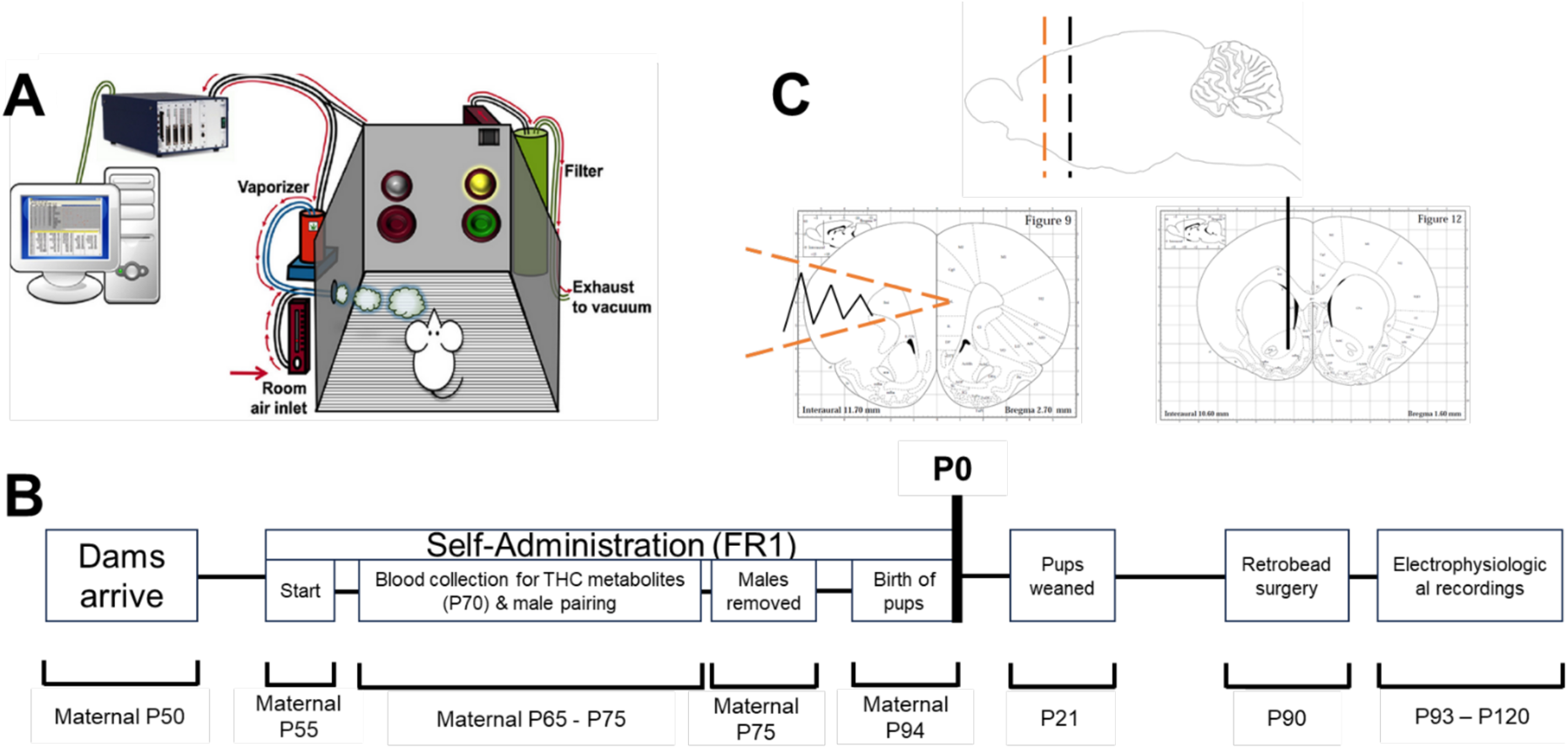
Experimental approach and timeline. **(A)** Rat dams acclimated to the laboratory environment for 5 days before beginning self-administration. Self-administration required a nosepoke in the active operandum, resulting in a 3-sec puff of vapor (THC or VEH) and a 1-min timeout. **(B)** After 10-days of self-administration, blood was collected for metabolite analyses and dams were paired with males for 10-days. Following day 10, dams were separated from males and continued self-administration until birth of pups (P0). Pups were weaned (P21) and maintained until P90. Offspring then underwent retrobead injection surgeries and electrophysiological recording. **(C)** Retrobeads were injected in the NAc (black bar) and incubated for approximately 3-days. Then offspring were euthanized, brain tissue collected and sliced, and neurons in the mPFC (orange bar) were recorded from.

The experimental timeline is provided in **Figure 1B**. Briefly, after 5 days of acclimation, rat dams were trained to nosepoke for cannabis vapor during twice daily, hour-long self-administration sessions in a vapor chamber room separate from the animal housing suite. The first session began two hours into the dark cycle and the second started two hours prior to the onset of the light cycle. The schedule of reinforcement was set to fixed ratio (FR)-1 such that each nosepoke in the active nosepoke not occurring during a timeout resulted in a vapor reinforcer. On the 10^th^ day of self-administration, female rats began a 10-day period of mating and then were single housed until the day of birth. Self-administration continued until 24-48 hr prior to birth. Pups were weaned on P21 and allowed to grow up in same-sex housing until they reached adulthood. Offspring were obtained from the maternal cannabis vapor cohort described in Weimar et al. (2023). Group sizes were: PCE (n=17; 10 M, 7 F), VEH (n=15; 5 M, 10 F).

### 2.4 Retrobead Surgery

Following maturation (P90), rats underwent retrobead injection surgery to label mPFC neurons that project directly to the NAc core. Rats were initially anesthetized via isoflurane (5%) in 100% oxygen in a sealed Plexiglas box. Once unconscious, rats were immobilized in a stereotaxic frame with anesthesia maintained at 2% isoflurane in 100% oxygen throughout surgery. Anesthesia was repeatedly tested via toe-pinch or tail-pinch. While immobilized, skin and fascia were removed on the skull and two craniotomies (<1 mm) were drilled. Using syringes rated for micro-injection (Hamilton, Reno, NV) fluorescent retrobeads (0.2 µl mg/mL) (Lumafluor, Durham, NC) were injected bilaterally into the NAc core (AP: +1.6; ML: ±1.8; DV: -6.8) (Paxinos & Watson, 1986) using a micro-injection pump (World Precision Instruments, Sarasota, FL) after a 2-min wait period once syringe tips reached the region of interest. After injection, incision sites were sutured, and rats were provided with post-surgery pain management (buprenorphine: 0.08 mg/kg; meloxicam: 1 mg/kg; enrofloxacin: 10 mg/kg) for 3 days. Neuronal labeling of the mPFC was verified for accuracy by examining retrobead placement in brain sections following electrophysiological recordings (**Figure 1C**).

### 2.5 Tissue Preparation and Ex Vivo Electrophysiology

Following retrobead injection and incubation (>3 days post-surgery), brain tissue was collected and whole-cell patch clamp electrophysiological recordings of fluorescently labeled neurons were conducted. Rats were deeply anesthetized with ketamine (100 mg/kg) and xylazine (7 mg/kg) and cardiac perfusions were performed with ice-cold artificial cerebrospinal fluid (aCSF) containing (mM): 147 Choline, 166 Na Ascorbate; 21 Na Pyruvate, 125 NaCl, 3 KCl, 1.2 KH_2_PO_4_, 1.2 MgSO_4_, 25 NaHCO_3_, 2 CaCl_2_, and 10 dextrose, bubbled with 95% O_2_ – 5% CO_2_ and brought to a pH of 7.40 using 1M HCl. Following perfusion, brain tissue was extracted and kept in ice-cold aCSF for the remainder of tissue collection. Tissue was placed in a vibratome (Leica VT1200S), horizontally mounted to a pedestal with cyanoacrylate glue, submerged in ice-cold aCSF, and coronal sections containing the mPFC were cut (400 µm thick). All tissue containing the mPFC was placed in a bath of warm (30°C) aCSF that was continuously bubbled with 95% O_2_ – 5% CO_2_ for 30 min. After 30 min, slices were removed from the 30°C bath and allowed to rest at room temperature (20°C) for 30 min. Following this rest period, recordings were performed on retrobead-containing mPFC neurons, as determined by fluorescence (excitation: 530 nm, emission: 590 nm). An upright Nikon FN1 microscope with a Nikon DS-Qi1Mc digital camera and NIS-elements AR imaging software were used for recordings. Initial recordings were completed with recording electrodes (3.0 – 4.5 MΩ) filled with a Cs-internal solution (mM): 10 CsCl, 130 Cs-Methanesulfonate, 11 EGTA, 1 CaCl_2_, 2 MgCl_2_, 10 HEPES, 2 Na_2_ATP, and 0.2 Na_2_GTP. Secondary experiments used a K+ internal solution to record changes in intrinsic membrane currents (mM): 6 NaCl, 4 NaOH, 130 K-gluconate, 11 EGTA, 1 CaCl_2_, 1 MgCl_2_, 10 HEPES, 2 Na_2_ATP, and 0.2 Na_2_GTP. For measurements of synaptic release, neurons were studied under voltage-clamp conditions with a MultiClamp 700A amplifier (Molecular Devices, Union City, CA) and held at V_H_ = -60 mV in whole-cell patch configuration. Only recordings with a series resistance of < 20 MΩ were used for experiments to ensure good access and maintenance of voltage clamp. Signals were filtered with a 1 kHz bezel filter and sampled at 20 kHz using Axon pClamp10 software (Molecular Devices).

The digitized waveforms of synaptic events were analyzed using an event detection and analysis program (MiniAnalysis, Synaptosoft, Decatur, GA) for all quantal synaptic currents. All events >10 pA were counted for frequency values. Fitting of quantal EPSC amplitudes and decay kinetics (90%–10%) were performed using a fitting protocol (MiniAnalysis) on >100 discrete events.

### 2.6 Statistics

For statistical comparisons, IBM SPSS Statistics (Version 27) was used. A two-way Analysis of Variance (ANOVA) was used to analyze the data, with condition (PCE, VEH) and sex (male, female) as between-subjects factors. Post-hoc analyses were conducted using Tukey’s HSD. Student’s t-tests were conducted as planned comparisons to test for treatment effects in each sex independently given our original hypotheses that were based on published data showing effects exclusively in male offspring (Bara et al., 2018). For all analyses, significance was defined by a *p* value < 0.05 and effect sizes (partial Eta squared (ƞp^2^) and Cohen’s *d* where appropriate) are provided for all statistical analyses.

## 3. RESULTS

### 3.1 Maternal Cannabis Vapor Self-Administration

We previously published that response-contingent delivery of vaporized cannabis extracts in pregnant rat dams supports stable rates of responding early in pregnancy (Weimar et al., 2023). This exposure resulted in physiologically active and pharmacologically relevant plasma THC concentrations and significantly lower birth weights in cannabis-exposed offspring relative to VEH-exposed offspring (Weimar et al., 2023), a finding that has also been reported in the human literature (Gunn et al., 2016). In this current study, we used offspring generated from the same cohort of dams used in Weimar et al., and as such, the self-administration data from these dams has been published previously (see Weimar et al., 2023). Male and female pups derived from either control or cannabis-exposed dams were compared as adults using whole-cell patch clamp electrophysiology in identified neurons from the mPFC. We analyzed the synaptic and neurophysiological properties of mPFC neurons labeled retrograde from the NAc core (see **Figure 1B** for experimental timeline). Injection sites in the NAc were verified and retrograde labeled neurons were brightly fluorescent in the mPFC (**Figure 1C**). For EPSC experiments, we recorded from N = 66 neurons from N = 32 rats (N = 9 Male/PCE, N = 6 Female/PCE, N = 9 Male/VEH and N = 8 Female/VEH) taken from N = 24 litters. For IPSC experiments, we recorded from N = 50 neurons from N = 23 rats (N = 7 Male/PCE, N = 4 Female/PCE, N = 6 Male/VEH and N = 6 Female/VEH) taken from N = 15 litters.

### 3.2 Impact of maternal cannabis on spontaneous glutamate release in identified NAc projecting mPFC neurons

To analyze changes in excitatory glutamatergic neurotransmission following maternal cannabis exposure, we recorded mPFC neurons labelled from the NAc in control (VEH) and cannabis exposed (PCE) male and female rats. Spontaneous excitatory post-synaptic currents (EPSCs), were recorded in labelled mPFC neurons held at V_H_ = -60 mV for a minimum of 15 min of recording time (**Figure 2A**). The frequency of spontaneous glutamate release varied across neurons and markedly between the sexes, which can be visualized in the cumulative probability graphs (**Figure 2B**). These plots show a difference in the rate of EPSCs between males and females under control conditions that is eliminated in PCE animals. Analysis of EPSC frequency was performed with a two-way ANOVA. Analyses revealed no main effect of PCE (*F*(1,62) = 0.23, *p* = 0.63, ƞp^2^ = 0.01) or sex (*F*(1,62) = 1.94, *p* = 0.17, ƞp^2^ = 0.04) on the frequency of spontaneous EPSCs, but there was a statistically significant interaction between PCE and sex (*F*(1,62) = 6.03, *p* = 0.017, ƞp^2^ = 0.09) (**Figure 2C**). Post-hoc analyses indicated no significant differences between PCE and VEH females though there was a trend for the frequency of EPSCs to be higher in PCE females relative to VEH females (*t*(20.7) = -1.95, *p* = 0.06, *d* = 0.78). It is worth noting that although we failed to reach the threshold for statistical significance, there was a medium-to large-sized effect (*d =* 0.78), which suggests that our inability to demonstrate a significanct effect was likely due to a lack of power. In contrast to females, there were no significant differences between PCE and VEH males, *t*(41) = 1.65, *p* = 0.11, *d* = 0.52 (**Figure 2C**). Examining EPSC frequency based on sex, regardless of condition, revealed that VEH females had significantly fewer EPSCs compared to VEH males, *t*(23.9) = - 2.95, *p* = 0.007, *d* = 1.0, and that PCE males and females had a similar frequency of EPSCs (*t*(36) = 0.78, *p* = 0.44, *d* = 0.28) (**Figure 2C**). These data collectively indicate that there are sex differences in the frequency of excitatory inputs to corticostriatal afferents in VEH offspring, which are not present following PCE.

**FIGURE 2:**
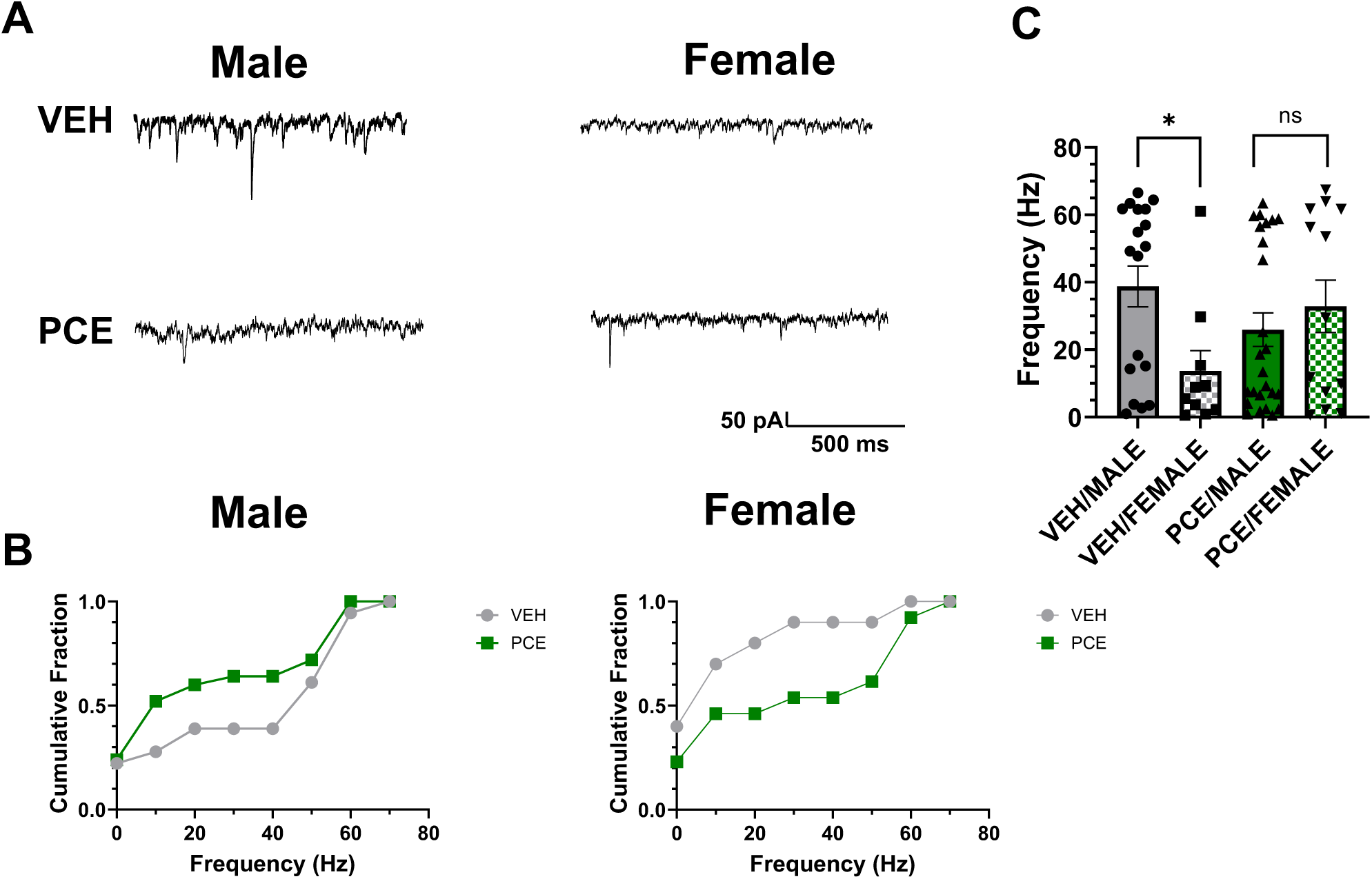
Prenatal cannabis exposure (PCE) abolishes sex differences in intrinsic excitability of corticostriatal efferents. **(A)** Representative current traces showing spontaneous EPSCs from labeled NAc projecting mPFC neurons. Recordings were divided between prenatal vehicle (VEH, top) and PCE (bottom) treated, as well as males (left) and females (right). **(B)** Cumulative fraction of EPSC recordings between VEH and PCE for males (left) and females (right) show categorically different frequency distributions between the male and female neurons as a function of treatment. **(C)** Analysis of EPSC frequencies demonstrated a significant sex difference in EPSC frequency that was not present in recordings taken from animals with PCE. Data are plotted as the mean ± stdev with the individual data points superimposed.

In addition to the frequency of glutamate release, the potential changes in postsynaptic glutamate receptor signaling were analyzed via EPSC waveform fitting for amplitude and decay-kinetics and compared via two-way ANOVA (**Figure 3**). We found that PCE significantly reduced the amplitude of spontaneous EPSCs (*F*(1,66) = 5.291, *p* = 0.025, ƞp^2^ = 0.08) independent of sex (*F*(1,62) = 0.13, *p* = 0.72, ƞp^2^ < 0.01) (**Figure 3A**), suggesting a change in postsynaptic receptor expression or location on the postsynaptic neuron. Analysis of the EPSC time constant (tau) as an indicator of glutamate receptor gating (closing and desensitization) showed no statistically significant differences based on PCE (*F*(1,62) = 1.18, *p* = 0.28, ƞp^2^ = 0.02) or sex (*F*(1,62) = 0.95, *p* = 0.33, ƞp^2^ = 0.02) and no significant interaction (*F*(1,66) = 0.83, *p* = 0.37, ƞp^2^ = 0.01) (**Figure 3B**).

**FIGURE 3:**
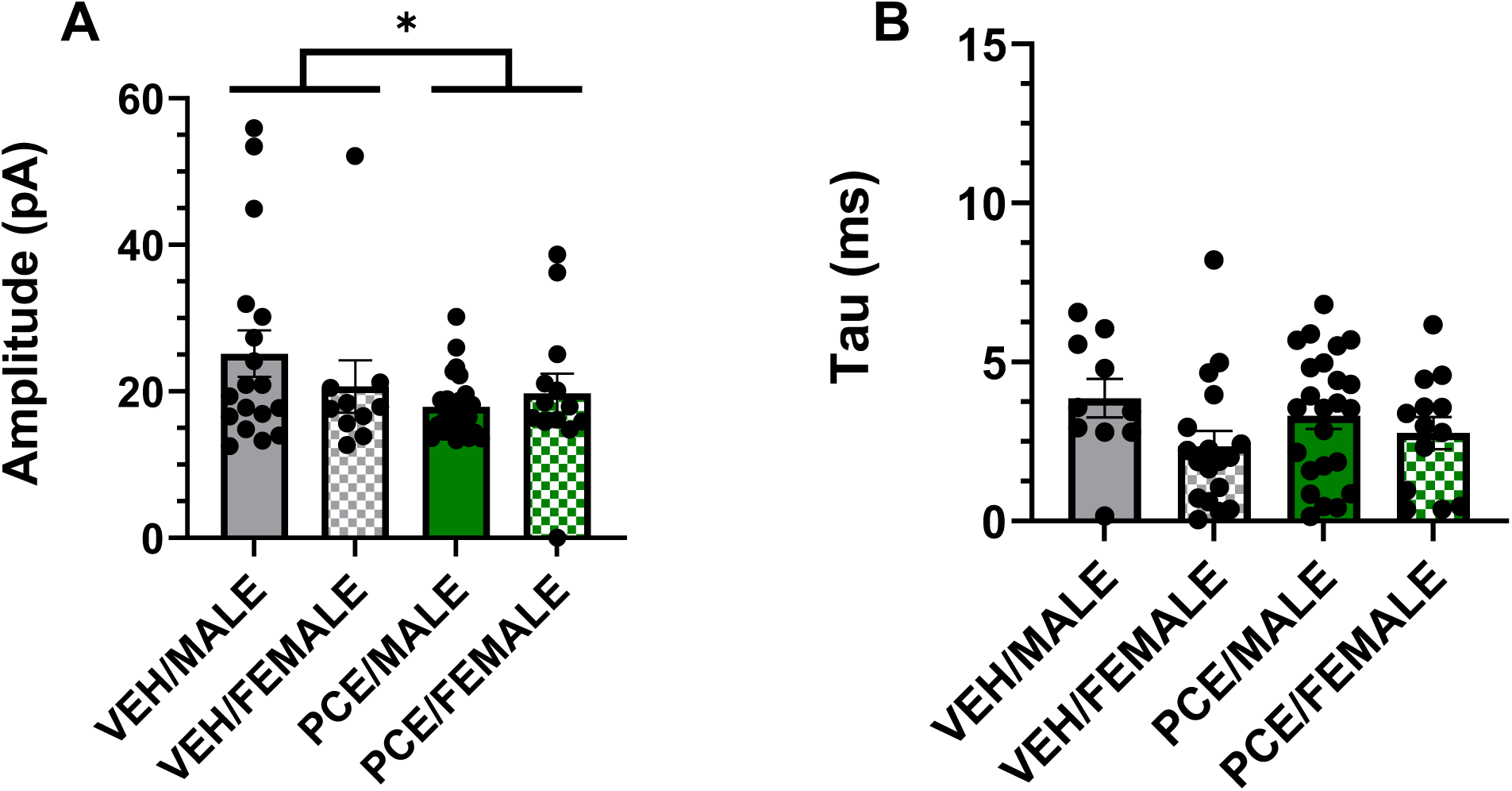
Prenatal cannabis exposure (PCE) reduces the amplitude, but not the decay kinetics, of spontaneous EPSCs in corticostriatal efferents of exposed offspring. **(A)** Spontaneous EPSCs analysis showed small, but statistically significant, reductions amplitude as a function of PCE with no effect of sex. **(B)** No significant differences in the decay kinetics (tau) of the EPSCs were observed based on prenatal treatment or sex. Data are plotted as the mean ± stdev with the individual data points superimposed.

### 3.3 Impact of maternal cannabis on inhibitory synaptic transmission of NAc projecting mPFC neurons

To isolate phasic inhibitory post synaptic currents (IPSCs), mPFC neurons were recorded at a holding potential of V_H_ = +20 mV for a minimum of 15 min to capture basal spontaneous IPSC frequency (**Figure 4**). At this holding potential the background potassium conductance produces additional noise in the holding current, but the upward deflecting IPSC waveforms remained clearly visible and analyzable (**Figure 4A**). Cumulative probability of IPSC frequency can be visualized in graphs (**Figure 4B**). In contrast with changes in glutamatergic signaling, we failed to detect a significant main effect of PCE (*F*(1,46) = 0.13, *p* = 0.72, ƞp^2^ < 0.01), sex (*F*(1,46) = 0.16, *p* = 0.70, ƞp^2^ < 0.03), or PCE x sex interaction (*F*(1,46) = 2.56, *p* = 0.12, ƞp^2^ = 0.05) on the frequency of spontaneous IPSCs (**Figure 4C**).

**FIGURE 4:**
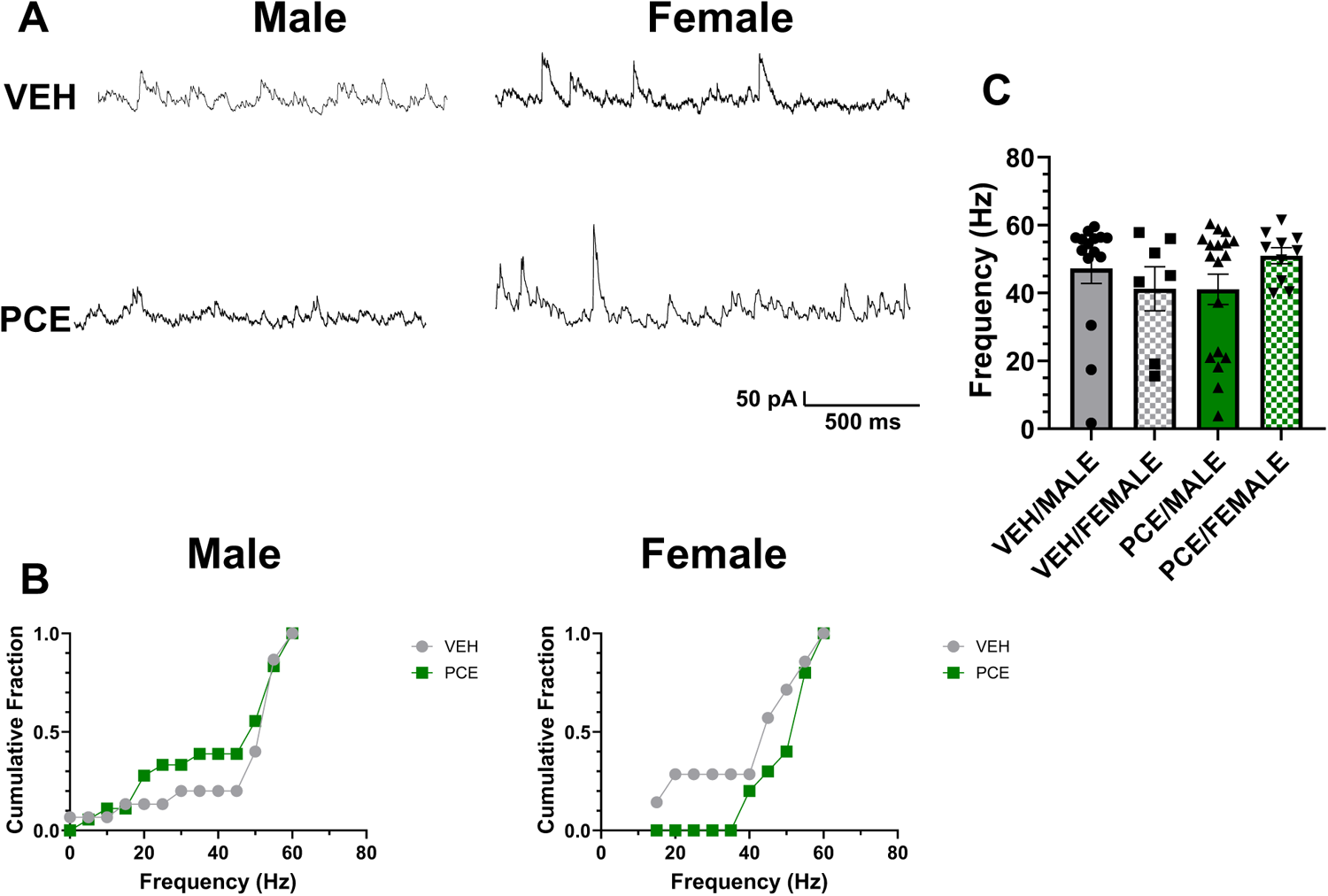
Prenatal cannabis exposure (PCE) does not alter the frequency of spontaneous IPSCs in corticostriatal efferent neurons of exposed offspring. **(A)** Representative current trace showing spontaneous IPSC traces from labeled NAc-projecting mPFC neurons. Neurons were held at +20 mV resulting in outward phasic IPSCs based on the chloride gradient. Recordings were divided between prenatal VEH (top) and PCE (bottom) treated, as well as males (left) and females (right). **(B)** Cumulative fraction of IPSC recordings between VEH and PCE for males and females as a function of treatment. **(C)** Analysis of IPSC frequencies for showed no statistical differences based on sex or prenatal exposure to cannabis. Data are plotted as the mean ± stdev with the individual data points superimposed.

### 3.4 Altered excitatory to inhibitory synaptic balance (EPSC / IPSC ratio) following prenatal cannabis exposure

The influence of synaptic transmission is largely determined by the balance of excitatory to inhibitory synaptic inputs on to a neuron over time. To capture this balance, we compared the EPSC / IPSC ratio of synaptic inputs on to labelled mPFC neurons (**Figure 5**). We found a statistically significant difference in the EPSC / IPSC ratio between male and female offspring (N = 7 – 15 neurons, *p* < 0.001, *d* = 1.82, T-test) in offspring from vehicle-treated dams that was not present following PCE (N = 10 – 18, *p* = 0.52, *d* = 0.04, T-test) (**Figure 5**). These differences, and PCE-induced changes, are driven by the marked changes in spontaneous glutamate release noted above.

**FIGURE 5:**
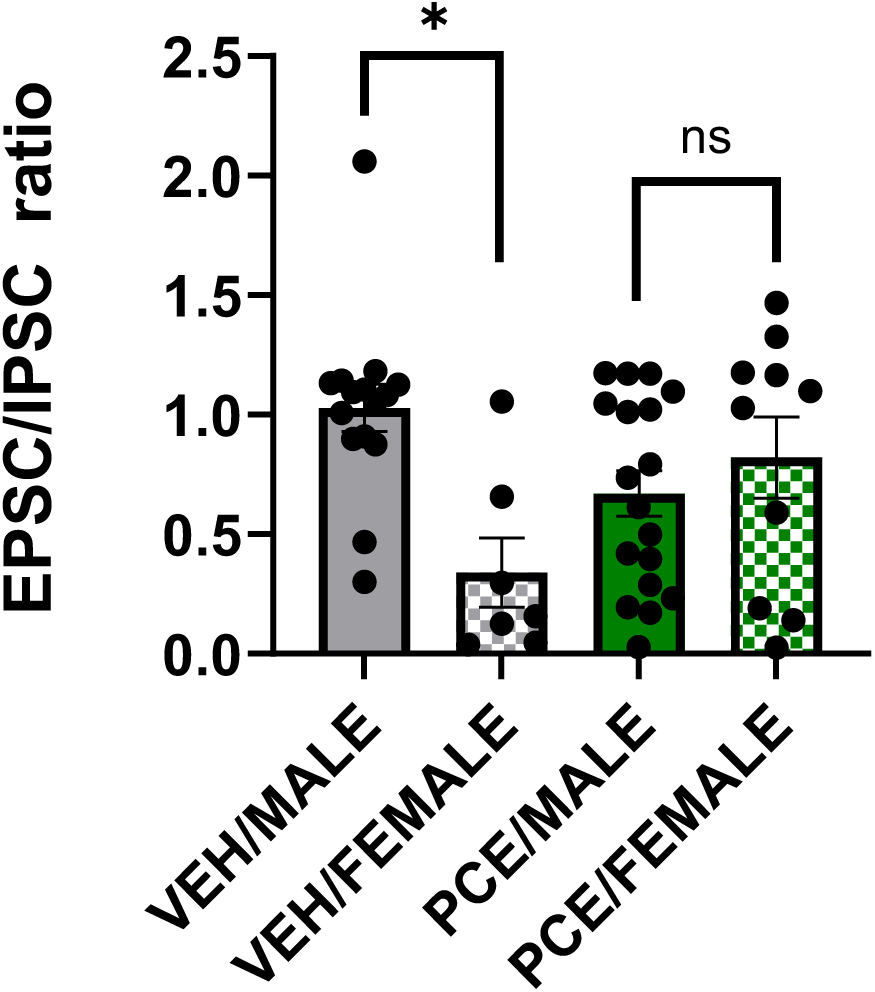
Prenatal cannabis exposure (PCE) abolishes sex differences in the EPSC / IPSC ratio of NAc projecting corticostriatal efferents. Under vehicle conditions there is a statistically significant difference in the EPSC / IPSC ratio on to labeled mPFC neurons from male and female rats. Prenatal cannabis treatment shifts the EPSC / IPSC ratio eliminating the sex difference observed in recorded mPFC neurons. The change in synaptic balance is driven by alterations in glutamatergic release as seen above. Summary data are plotted as the mean ± stdev with the individual data points superimposed.

### 3.5 Maternal cannabis produced no change on intrinsic membrane currents of identified NAc projecting mPFC neurons

Neurophysiological changes due to PCE may also alter intrinsic membrane currents in mPFC neurons (**Figure 6**). To test this possibility, we analyzed the intrinsic membrane conductance of labelled mPFC neurons using voltage steps (starting at V_H_ = -100 mV, increasing 10 mV per step, ending at V_H_ = +80 mV) and producing current-voltage (IV) plots (**Figure 6A and C**). We found that PCE failed to significantly alter membrane conductance at resting membrane potential of -60 mV across treatment *(F*(1,56) = 0.094, *p* = 0.76, ƞp^2^ = 0.04) and between sex (*F*(1,56) = 0.023, *p* = 0.880, ƞp^2^ = 0.07) (**Figure 6B and D, left panels**). Two-way ANOVA of cell current at a membrane voltage of +60 mV showed no main effects of sex (*F*(1,57) = 0.658, *p* = .421, ƞp^2^ = 0.02), PCE (*F*(1,57) = 1.072, *p* = .305, ƞp^2^ = 0.01), nor an interaction between PCE and sex (*F*(1,57) = 0.016, *p* = .901, ƞp^2^ < 0.01) (**Figure 6B & D, right panels**).

**FIGURE 6:**
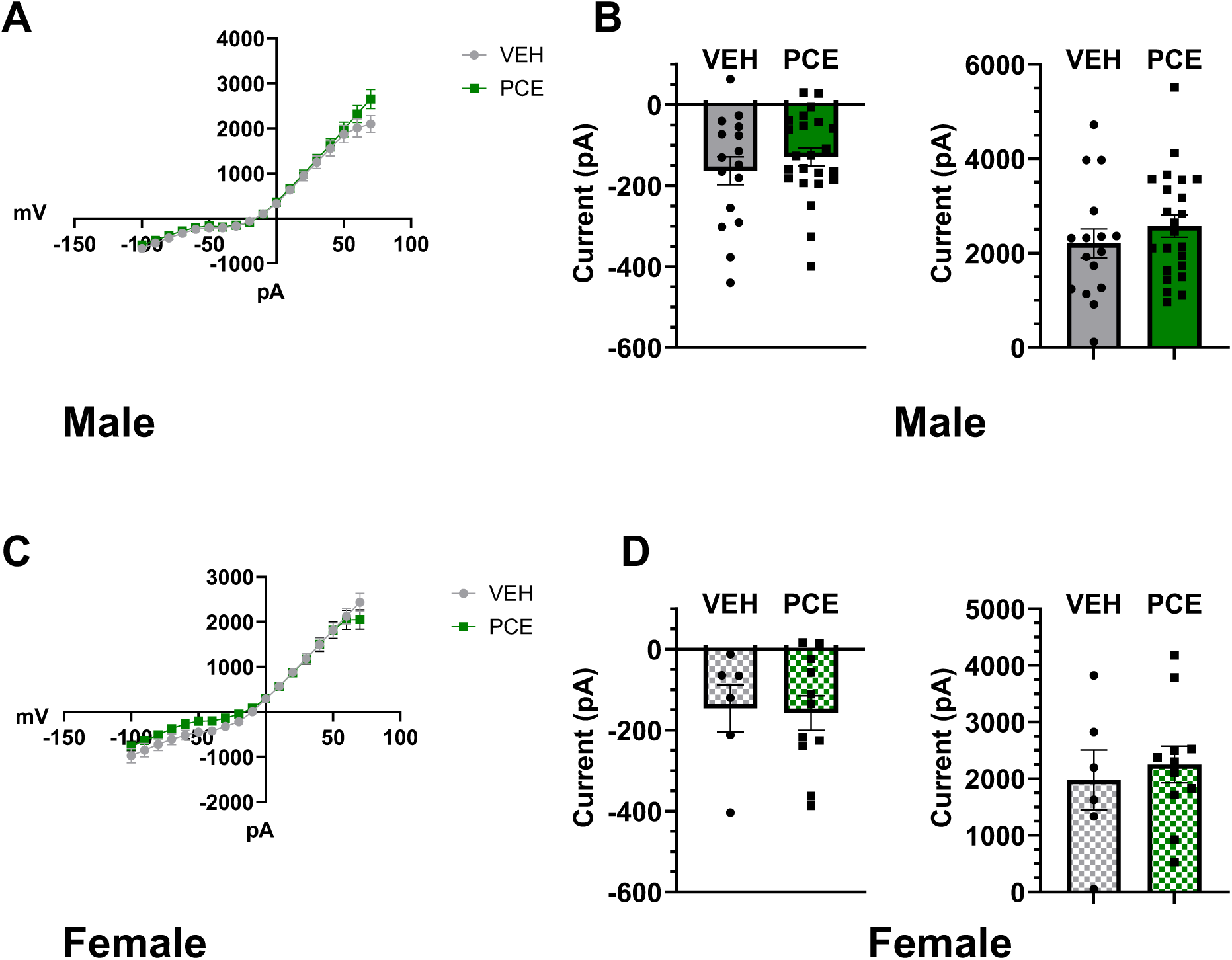
Prenatal cannabis exposure fails to alter intrinsic membrane conductances. Average current/voltage curves from NAc projecting mPFC neurons of VEH and PCE males. **(A)** and females **(C)**. Data are plotted as the mean ± stdev current (pA) across voltage steps. Analysis of membrane currents from males **(B)** and female **(D)** mPFC neurons at approximately resting potential (-60 mV, left panels) and at a depolarized potential (+60 mV, right panels) revealed no statistical differences based on prenatal cannabis treatment. Summary data are plotted as the mean ± stdev at each potential with individual data points superimposed.

## 4. DISCUSSION

The current set of experiments report mPFC neurons projecting to the NAc maintain sex specific differences in excitatory to inhibitory synaptic balance (glutamate to GABA) that is lost following PCE. VEH-exposed males showed a greater frequency of spontaneous glutamate release compared to VEH-exposed females (**Figure 2**) with no differences in spontaneous GABA release (**Figure 4**), resulting in a greater excitatory to inhibitory tone in males specifically (**Figure 5**). Treatment of the pregnant dams with vaporized cannabis (PCE, see Methods and **Figure 1** for protocol) increased the frequency of spontaneous glutamate release selectively in females and eliminated the sex difference in excitatory to inhibitory tone suggesting sex-specific changes in the presynaptic release mechanisms of glutamate. In contrast, PCE decreased the amplitude of glutamatergic EPSCs in both sexes suggesting a common change in postsynaptic receptor expression or trafficking. Finally, we did not observe any treatment effect or sex difference in the intrinsic membrane properties of labeled mPFC neurons (**Figure 6**). Together, these results show a subtle and nuanced change in synaptic processing in PCE males that may impact mPFC firing and contribute to observed behavioral and learning changes mediated by corticostriatal circuitry.

Compared to the previous literature, the present findings highlight a few key concepts for consideration. First, our findings partially align with those of Scheyer et al. (2020a) who reported that low injected current resulted in fewer action potentials in rats that experienced perinatal THC exposure. The data here similarly show that amplitude in PCE rats was reduced, which is indicative of a reduced ability for excitatory input to depolarize the cell. Taken together with the Scheyer et al. (2020a) findings, our results indicate that both PCE and perinatal THC exposure impact the threshold for depolarization, reducing mPFC firing activity in animals that are cannabinoid-exposed at these developmental time points. This is not true in the case of perinatal WIN exposure (Scheyer et al., 2020b), in which rats that were exposed to WIN from P1 to P10 showed no changes in neural firing following injected current. This difference may be due to the use of different cannabinoids in these studies (phytocannabinoids vs. WIN). For instance, individual cannabinoids such as WIN and THC have a very different chemical structure (Compton et al., 1992) and display functional selectivity at CB1Rs, leading to activation of distinct signaling pathways (Laprairie et al., 2014). Additionally, administration of WIN leads to rapid CB1R internalization, whereas THC causes little receptor internalization (Hsieh et al., 1999). Furthermore, THC and other phytocannabinoids interact with other receptor types (*e.g.* 5-HT3, TRPV, µ & δ opioid receptors), whereas WIN is a highly selective CB1R agonist (Compton et al., 1992). These differences may contribute to the different findings of Scheyer et al. (2020b) versus those in Scheyer et al. (2020a) and the present study. The difference between prenatal WIN exposure and THC is further supported by data from Bara et al. (2018). Male rats exposed to prenatal THC/WIN showed changes in rheobase, whereas females were spared (Bara et al., 2018). This indicates that there may be sex-specific changes following prenatal THC/WIN exposure, but our model of maternal cannabis use did not replicate this. The findings here indicate both males and females show reductions to ESPC amplitude, highlighting that the amplitude of excitatory currents in NAc-projecting mPFC neurons in both sexes are similarly impacted by PCE.

Most recent studies examining effects of PCE have noted sex differences, with alterations typically observed in exposed male offspring (Bara et al., 2018; Frau et al., 2019; Sagheddu, et al., 2021; Frau and Melis, 2023). These results directly contrast with our data showing a medium-to large-sized effect (*d =* 0.78) of PCE on the frequency of EPSCs in female, but not male, rats. Although we failed to reach the threshold for statistical significance, this was likely due to a lack of power rather than a lack of effect per se. It is worth noting that we are the first group to our knowledge to interrogate effects of PCE on a specific subpopulation of NAc-projecting mPFC neurons. As such, a likely explanation for discrepancies between our data and published data is that PCE has effects on this subpopulation of mPFC neurons in females that are masked when utilizing more unbiased cell sampling approaches. The mPFC exerts top-down control over a number of limbic and midbrain regions including the thalamus, basolateral amygdala, ventral tegmental area, dorsal raphe nucleus, lateral habenula, and periaqueductal gray, among others, which each participate in optimizing higher-order cognitive functioning (see Anastasiades & Carter, 2021 for review). Future studies comparing effects of PCE on excitatory and inhibitory transmission in different mPFC projection neurons are needed to fully understand the potential sex-specific effects of PCE and their behavioral consequences.

Another methodological disparity that could have resulted in differences between our results and those of others is the method of drug administration. Our study uses a self-administration approach involving response-contingent delivery of vaporized cannabis extracts, whereas the studies described above used an injection protocol involving THC or WIN. As reported by Baglot et al. (2022), metabolism of THC occurs at different rates depending on route of administration, such that cannabinoid concentration reaching the developing fetus differs based on administration route. Although our maternal self-administration method results in lower fetal cannabinoid exposure compared to previous work, we still show alterations that are directly related to PCE, indicating that even low-dose cannabis exposure during gestation can alter brain function in adulthood. Pulmonary administration (*i.e.* vaping, smoking) is the preferred method of cannabis use in humans (Schauer et al., 2020), which our model mimics. Thus, our data indicate that even relatively low levels of cannabis exposure during pregnancy are capable of producing long-term neurobiological effects in exposed offspring.

The increase in action potential probability in corticostriatal neurons may help partially explain the diminished impulse control (Fried et al., 1998) found in humans exposed to cannabis *in utero*. Local field potential (LFP) recordings in rats performing a 5-Choice Serial Reaction Time Task (5-CSRTT) allowed for the categorization of rats as highly impulsive or non-impulsive, based on premature responses and magnitude of LFP signals between the mPFC and NAc (Donnelly et al., 2014). Higher amplitude LFPs that were not phase-amplitude coupled were shown to be correlated with highly impulsive, premature responding. In contrast, non-impulsive rats showed lower amplitude LFPs that were phase-amplitude coupled between the mPFC and nAc (Donnelly et al., 2014). This desynchrony between the mPFC and NAc after PCE may be explained by altered excitability of NAc-projecting mPFC neurons. Disconnection of the mPFC to NAc by muscimol was shown to increase premature responding in rats tested in a 5-CSRTT (Feja & Koch, 2014). These studies highlight the importance of mPFC-to-NAc communication for regulating impulsive behavior. Our findings further show that a decrease in EPSC amplitude in NAc-projecting mPFC neurons following PCE may contribute to the lack of impulse control observed in human longitudinal studies of PCE (Fried et al., 1998). PCE likely drives aberrant communication between mPFC and NAc, causing high amplitude LFPs in both the mPFC and NAc that are not phase-amplitude coupled, increasing impulsive behavior in exposed offspring.

Interestingly, two distinct populations of corticostriatal projection neurons were found during recording in both conditions and sexes: those with a high-frequency of activity, >40 Hz, and those with a low-frequency of activity, <25 Hz. These populations of neurons may be separate types of projection neurons, modulating activity in the NAc in different manners entirely. Additional recordings that further separate neuron types may provide further insight as to how the mPFC controls the NAc, as well as how PCE impacts these two unique subpopulations of corticostriatal neurons. Finally, the mPFC consists of two subregions that project to the NAc, the infralimbic and prelimbic cortices. These two regions have been shown to differentially impact multiple facets of cognition, such as attention and impulsivity (see Vertes 2004 for review). Recording from populations of projection neurons in these specific subregions may reveal further mechanistic understanding of the impact of PCE on these specific aspects of cognition.

There are clear sex differences in the development of the rodent mPFC (Premachandran et al., 2020). Our findings add to this body of literature by showing that adult female rats exhibit significantly fewer spontaneous EPSCs in NAc-projecting mPFC neurons compared to males in the control condition. Less excitatory drive onto these neurons in females could help to explain the propensity for risk aversion in females that has been reliably shown using rodent decision-making tasks that involve the mPFC-to-NAc pathway (see Orsini et al., 2022 for review). Although we did not measure stage of estrous in this study, others have shown that sex hormones also modulate the structure and function of mPFC neurons. For instance, ovariectomy leads to dendritic spine loss in the mPFC, which is reversed by estrogen administration (Luine et al., 2006; Shansky et al., 2010). Furthermore, estrogen increases dendritic spines in infralimbic (IL) mPFC neurons that project to the basolateral amygdala in adult rats (Shansky et al., 2010). At a functional level, female mice in proestrus exhibit a decrease in spontaneous EPSCs compared to mice in diestrus when recording from pyramidal neurons in layer 5 of the infralimbic cortex (IL; Galvin & Ninan, 2014). Remarkably, estrogen-induced changes in spine density and functional activity often coincide with enhanced performance in mPFC-dependent behavioral tasks (Inagaki et al., 2012; Velazquez-Zamora et al., 2012; Luine and Frankfurt, 2013). While our recordings are from mPFC neurons in the PL as opposed to IL, these studies suggest that the organization and function of mPFC neurons is both sex and hormone dependent, and that these factors may similarly contribute to our findings. Future studies will be required to fully dissect the impact of sex and hormones/estrus stage on spontaneous (and evoked) activity in NAc-projecting mPFC neurons, as well as their contribution to facets of risky decision making and cognitive flexibility that are known to require this neuronal population.

In conclusion, the current study shows that, in NAc-projecting mPFC neurons, there are sex differences in ESPC frequency, with females showing fewer EPSCs compared to males. VEH males and females show distinct differences that are not apparent in PCE males and females. Finally, PCE decreases EPSC amplitude irrespective of sex, which results in reduced ability for excitatory input to depolarize the cell, thereby decreasing excitability of these corticostriatal neurons. These findings may help to explain differences seen in human populations regarding attentional deficits and the increase in impulsive behaviors following PCE.

## ACKNOWLEDGEMENTS

This study was supported by funds provided for medical and biological research by the State of Washington Initiative Measure No. 171 (RJM, HVW, and DEG). The authors would also like to thank Hayden Wright, Nicholas Glodosky, Alexandra Malena, and Ginny Park for their assistance with animal care. The authors have no conflicts of interest. These contents do not represent the views of the U.S. Department of Veterans Affairs or the United States Government.

